# Conserved regulation of omptin proteases by the PhoPQ two-component regulatory system in Enterobacteriaceae

**DOI:** 10.1101/2020.02.28.965145

**Authors:** Youn Hee Cho, Monir Riasad Fadle Aziz, Tanuja Sutradhar, Jasika Bashal, Veronica Cojocari, Joseph B. McPhee

**Author notes:** These authors contributed equally to the work (Youn Hee Cho, Monir Riasad Fadle Aziz).

## Abstract

Bacteria that colonize eukaryotic surfaces interact with numerous host-produced molecules that have antimicrobial activity. Bacteria have evolved numerous strategies to both detect and resist these molecules, and in gram-negative bacteria these include alterations of the cell surface lipopolysaccharide structure and/or charge and the production of proteases that can degrade these antimicrobial molecules. Many of the lipopolysaccharide alterations found in enteric bacteria are controlled by the PhoPQ and PmrAB two-component regulatory systems. Here, we show that omptin family proteases from *Escherichia coli* and *Citrobacter rodentium* are induced by growth in low Mg^2+^. We further show that deletion of PhoP eliminates omptin protease activity, transcriptional regulation and protein levels. We identify conserved PhoP-binding sites in the promoters of the *E. coli* omptin genes, *ompT*, *ompP* and *arlC* as well as in *croP* of *Citrobacter rodentium* and show that mutation of the putative PhoP-binding site in the *ompT* promoter abrogates PhoP-dependent expression. Finally, we show that despite the conserved PhoP-dependent regulation, each of the *E. coli* omptin proteins has differential activity toward a particular substrate, suggesting that each omptin may contribute to resistance to a particular repertoire of host-defense peptides, depending on the particular environment in which each evolved.

## Introduction

Enterobacteriaceae are a family of gram-negative bacteria that, in addition to serving as commensals in the mammalian gastrointestinal tract, are also important pathogens. *Escherichia coli* is a common cause of acute gastroenteritis as well as extraintestinal infections, including urinary tract infections and sepsis (Croxen and Finlay, 2010; Croxen *et al.*, 2013). Bacteria that colonize or cause disease in the human gastrointestinal tract or other mucosal surfaces encounter a significant number of environmental stressors and they have evolved sophisticated mechanisms for both detecting these stressors and for mounting an appropriate defensive response. These bacterial defense systems are critical for the lifestyle of the microbe and understanding how these systems are regulated and how they function represents an important research goal.

In order to protect themselves, mammalian hosts have developed a complex and sophisticated system of both innate and adaptive immune defences. The innate immune system is the first line of defense and includes two components: cellular components like neutrophils and macrophages as well as humorally produced molecules with direct antimicrobial activity and innate pattern recognition. These soluble components of the innate immune response include cationic antimicrobial peptides, secreted pattern recognition receptors and components of the complement signaling cascade. The complement pathway is a fairly complex component of the innate immune response (Skattum *et al.*, 2011). In it, soluble serum components of the host recognize conserved structures on the surfaces of microbial cells, leading to the formation of a membrane attack complex and bacterial lysis. The pathway also serves to recruit cellular innate immune components, thereby leading to enhanced infection control. In spite of this, numerous bacteria have evolved mechanisms to resist the activity of complement, in order to enhance virulence (Potempa and Potempa, 2012).

Cationic antimicrobial peptides are a structurally diverse group of molecules that contribute to both constitutive and inducible host defense (Haney *et al.*, 2017). These peptides are able to bind to and penetrate bacterial membranes, leading to localized aggregation, reorganization and disruption (Aquila *et al.*, 2013). This membrane activity is thought to be the main mechanism by which the majority of cationic antimicrobial peptides function. In humans, these molecules are broadly grouped by structure into α-helical peptides as well as those with β-sheet structure. In the former group, the most important peptide is LL-37. This peptide is a cathelin-family peptide and is produced by neutrophils, macrophages as well as by mucosal epithelial cells throughout the body (Zanetti, 2004). During inflammation, it is upregulated in epithelial cells as well as secreted at higher levels by infiltrating cells of the innate immune system (Kusaka *et al.*, 2018). LL-37 has broad immunomodulatory effects as well as direct antibacterial effects, principally by interactions with the bacterial cell envelope, leading to loss of cellular integrity and cell death (Gudmundsson *et al.*, 1996; Turner *et al.*, 1998; Bowdish *et al.*, 2006). Together, the complement pathway and cationic antimicrobial peptides protect mucosal and endothelial surfaces from potential microbial attack.

Bacteria are not willing participants in this host-mediated defense system, however. In the Enterobacteriaceae, many species have evolved regulatory protein systems that respond directly to environmental cues, including changes in external osmolarity or pH, exposure to host or bacterial derived signaling molecules and to the presence of host-defense peptides themselves. In *E. coli*, the best characterized system for resistance to host-defense peptides involves two separate two-component signaling pathways, the PhoPQ and PmrAB systems. The net effect of activating the PhoPQ signaling system is that the bacteria become resistant to cationic host-defense peptides due to reduced peptide binding and altered membrane hydrophobicity.

In addition to reducing surface peptide binding, bacteria also express outer membrane-associated proteases that can degrade proteins that are in or on the bacterial surface (Sugimura and Nishihara, 1988). These omptin family proteins, named for the prototypical OmpT protein, are 10-stranded β-barrels in which the protease catalytic site is found in the surface-associated loops (Vandeputte-Rutten *et al.*, 2001). Numerous OmpT homologs are found throughout the Enterobacteriaceae, including PgtE in *Salmonella*, CroP in *Citrobacter*, OmpP and ArlC in *E. coli*, SopA in *Shigella*, and Pla of *Yersinia pestis* (Johanna Haiko *et al.*, 2009). These proteins can cleave some host-defense peptides and they contribute to resistance to this class of molecules in numerous bacterial species. Here, we wanted to determine whether omptin-family proteins are regulated by the PhoPQ system as well as investigate whether different omptins exhibit altered substrate specificity.

## Materials and methods

### Bacterial strains, genetic manipulations and growth conditions

All strains and plasmids used in this study are shown in table 1. Bacteria were routinely grown in lysogeny broth for culturing and for molecular manipulations. Chemically competent cells (*E. coli* and *C. rodentium*) were prepared as previously described (Green and Rogers, 2013). For antibiotic selection, we routinely used ampicillin at 100 μg/ml or kanamycin at 50 μg/ml. For defined media composition we used a modified N-minimal medium (Nelson and Kennedy, 1971). Briefly, media contained 5 mM KCl, 7.5 mM (NH_4_)_2_SO_4_, 0.5 mM K_2_SO_4_, 1 mM KH_2_PO_4_, 0.1 mM Tris-HCl pH 7.4, 10 mM or 10 μM MgCl_2_, and 0.2% glucose.

**Table 1:**
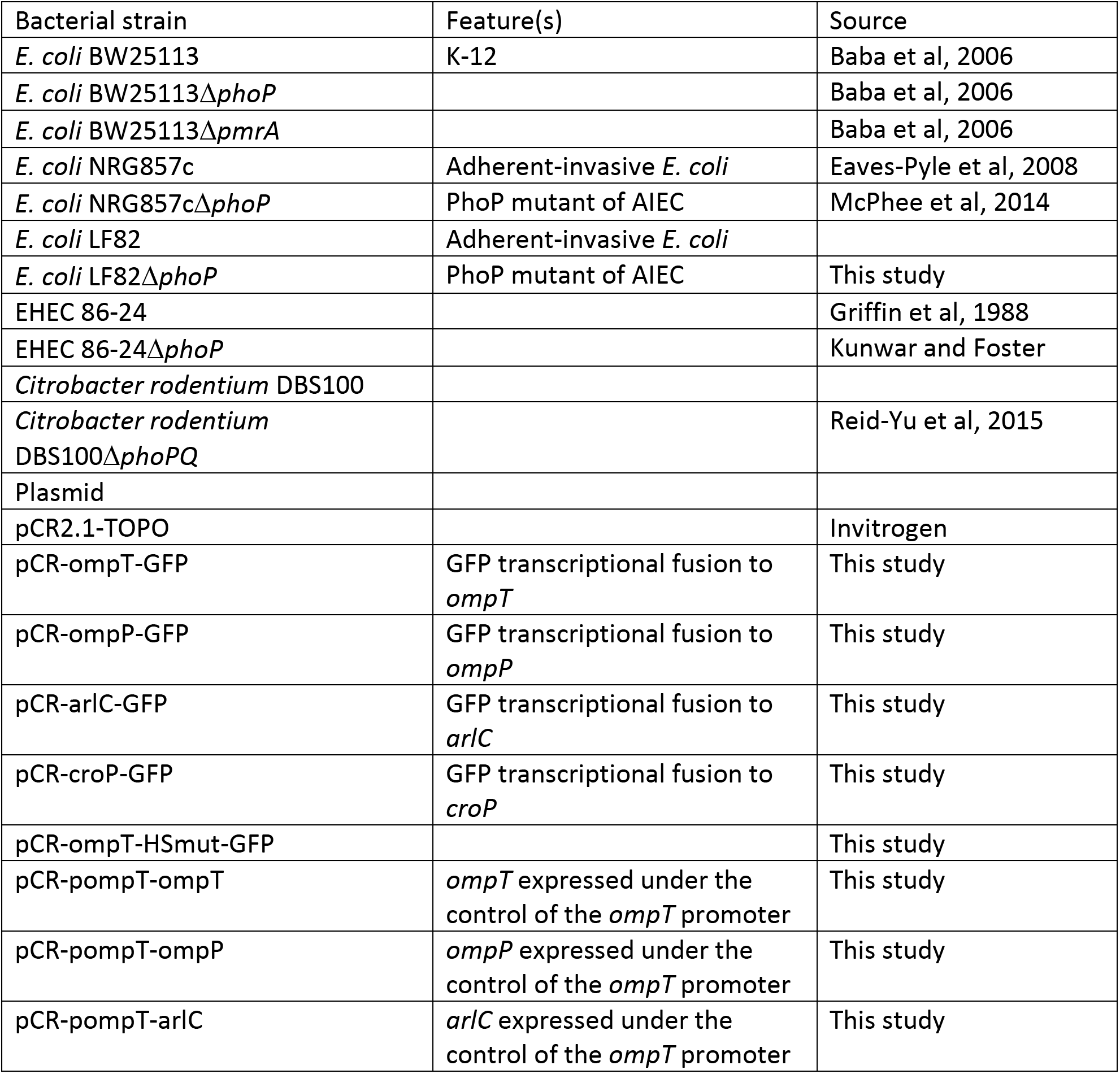
Strains and plasmids used in this study

| Bacterial strain | Feature(s) | Source |
| --- | --- | --- |
| *E. coli* BW25113 | K-12 | Baba et al, 2006 |
| *E. coli* BW25113Δ*phoP* | | Baba et al, 2006 |
| *E. coli* BW25113Δ*pmrA* | | Baba et al, 2006 |
| *E. coli* NRG857c | Adherent-invasive *E. coli* | Eaves-Pyle et al, 2008 |
| *E. coli* NRG857cΔ*phoP* | PhoP mutant of AIEC | McPhee et al, 2014 |
| *E. coli* LF82 | Adherent-invasive *E. coli* |  |
| *E. coli* LF82Δ*phoP* | PhoP mutant of AIEC | This study |
| EHEC 86-24 |  | Griffin et al, 1988 |
| EHEC 86-24Δ*phoP* | | Kunwar and Foster |
| *Citrobacter rodentium* DBS100 |  |  |
| *Citrobacter rodentium* DBS100Δ*phoPQ* | | Reid-Yu et al, 2015 |
| Plasmid |  |  |
| pCR2.1-TOPO |  | Invitrogen |
| pCR-ompT-GFP | GFP transcriptional fusion to *ompT* | This study |
| pCR-ompP-GFP | GFP transcriptional fusion to *ompP* | This study |
| pCR-arlC-GFP | GFP transcriptional fusion to *arlC* | This study |
| pCR-croP-GFP | GFP transcriptional fusion to *croP* | This study |
| pCR-ompT-HSmut-GFP |  | This study |
| pCR-pompT-ompT | *ompT* expressed under the control of the *ompT* promoter | This study |
| pCR-pompT-ompP | *ompP* expressed under the control of the *ompT* promoter | This study |
| pCR-pompT-arlC | *arlC* expressed under the control of the *ompT* promoter | This study |

### FRET based omptin protease assay

We developed a probe in which an internal 11 amino acid sequence of the human peptide LL-37 (LLGDFFRKSKEKIGKEFKRIVQRIKDFLRNLVPRTES) were labeled on the N-terminus with a fluorescent probe while the C-terminus was tagged with a proprietary quenching molecule (5-FAM-GKEFKRIVQRI-K(QXL520)). When this molecule is cleaved, quenching is relieved and fluorescence can be monitored. Strains were grown overnight in LB or N-minimal medium at 37°C with shaking. The overnight culture was washed, diluted 1:50 in fresh LB or N-minimal medium and grown to mid-log phase at 37°C. The cultures were washed twice in 10 mM HEPES buffer pH 7.2, then normalized to an OD_600_ of 0.5. The FRET substrate was added to a final concentration of 7 μg/mL to ~1.1 × 10^7^ cells or HEPES buffer as a blank. Fluorescence was measured over 1 hour at 25°C with an excitation wavelength of 485nm and emission wavelength of 528nm. The initial slope of the activity was calculated and the measurements shown represent the average of at least three independent experiments.

### Transcriptional reporter assay

We carried out splicing by overlap extension polymerase chain reaction (SOE-PCR) to create transcriptional reporter fusion between the promoters of *ompT*, *ompP*, *arlC*, *croP*, or *pgtE* and the GFP-mut3 gene. To do this, we used purified genomic DNA from strains containing these alleles and for the GFP-*mut3* PCRs, we used purified template DNA from p3174 (Gerlach *et al.*, 2009). The promoters were amplified using P1 and P2 while the GFP gene was amplified using P3 and P4. The purified products were then joined by PCR using P1 and P4 along with the purified products of the promoter and GFP reactions (~100 ng each product in a 25 µl reaction made according to the manufacturer’s instructions).

The spliced DNA fragments were then cloned to pCR2.1-TOPO according to the manufacturer’s instructions and the construct was sequence verified by Sanger sequencing. For the construction of the pCR-ompT-HSmut-GFP PhoP-binding site mutant plasmid, we carried out site direct mutagenesis of pCR-ompT-GFP according to the Quikchange II site-directed mutagenesis kit (Agilent Technologies).

The plasmids were transformed to the appropriate strain using electroporation or rubidium chloride mediated chemically competent cell treatment as appropriate. For carrying out the transcriptional reporter assays, overnight cultures were grown in LB + 20 mM MgSO4. The next day, the cultures were washed 2×in 1×N-minimal salts and resuspended to and OD600 of ~0.02 in N-minimal medium containing either 10 mM MgSO4 and 0.0015% FeSO4 (high Mg, low Fe), 10 μM MgSO_4_ and 0.0015% FeSO4 (low Mg, low Fe) or 10 mM MgSO4 and 0.15% FeSO4 (high Mg2+, high Fe3+). The inoculated cultures were then pipetted into black sided, clear-bottomed 96 well culture plates and measured every 30 minutes for fluorescence with excitation at 485 nm and emission at 530 nm along with for absorbance at 600 nm. The ratio of fluorescence to absorbance was calculated at a mid-logarithmic growth point and they are reported as relative fluorescence unit/OD600.

### Western blot for OmpT and DnaK

Bacteria were grown with shaking in LB at 37°C overnight. Cells were then separately subcultured in N-minimal high Mg^2+^ media, N-minimal low Mg^2+^ media, and N-minimal high Mg^2+^ high Fe^3+^ media with shaking at 37 °C until early stationary phase was reached. Cells obtained were normalized to OD_600_ of 1 (2 × 109 cells) before pellet collection via centrifugation at 17,000 × G for 2 minutes and the supernatant was discarded. The pellets were resuspended in pre-chilled sterile PBS, washed once and stored at −20°C overnight. Next, the pellets were resuspended in 1 × DNAse I reaction buffer (10 mM Tris-HCl, 50 mM NaCl, and 10 mM MgCl_2_ at pH 7.5) and subjected to lysis via three freeze-thaw cycles prior to incubation with 10 mg/mL DNAse I at 37 °C for 1 hour. Samples were incubated at 100°C for 10 minutes in 1 × BCA compatible lysing buffer (4 % SDS and 0.2 M Tris-HCl at pH 6.8). Total protein concentration in each sample was measured using Pierce^®^ BCA Assay Kit (Thermo Fisher Inc.), and samples were mixed with post-lysis Laemmli buffer (2 % SDS, 60 % glycerol, 1.2 M DTT, and 0.1 M Tris-HCl at pH 6.8) before storage at −20 °C. The protocol was reproduced with BW25113, BW25113ΔpmrA and BW25113∆*phoP* strains in N-minimal low Mg^2+^ media.

Total protein concentrations in samples were normalized to 1600 µg/mL before loading. Proteins were separated on a 12% Western blot for OmpT and DnaK. Bacteria were grown with shaking in LB at 37°C overnight. Cells were then separately subcultured in N-minimal high Mg^2+^ media, N-minimal low Mg^2+^ media, and N-minimal high Mg^2+^ high Fe^3+^ media with shaking at 37 °C until early stationary phase was reached. Cells obtained were normalized to OD_600_ of 1 (2 × 109 cells) before pellet collection via centrifugation at 17,000 × G for 2 minutes and the supernatant was discarded. The pellets were resuspended in pre-chilled sterile PBS, washed once and stored at −20°C overnight. Next, the pellets were resuspended in 1 × DNAse I reaction buffer (10 mM Tris-HCl, 50 mM NaCl, and 10 mM MgCl_2_ at pH 7.5) and subjected to lysis via three freeze-thaw cycles prior to incubation with 10 mg/mL DNAse I at 37 °C for 1 hour. Samples were incubated at 100°C for 10 minutes in 1 × BCA compatible lysing buffer (4 % SDS and 0.2 M Tris-HCl at pH 6.8). Total protein concentration in each sample was measured using Pierce^®^ BCA Assay Kit (Thermo Fisher Inc.), and samples were mixed with post-lysis Laemmli buffer (2 % SDS, 60 % glycerol, 1.2 M DTT, and 0.1 M Tris-HCl at pH 6.8) before storage at −20 °C. The protocol was reproduced with BW25113, BW25113ΔpmrA and BW25113∆*phoP* strains in N-minimal low Mg^2+^ media. Proteins were separated on 12% SDS-polyacrylamide gels in 1 × SDS-PAGE running buffer (25 mM Tris-HCl pH 8.3, 200 mM glycine, 0.1 % SDS), followed by electroblotting to nitrocellulose membranes at 200 V for 1 hour in 1X transfer buffer (10 % 1 × Tris-glycine pH 8.3 and 20 % methanol). Membranes were blocked with 5 % skim milk in TBST (20 mM Tris-HCl pH 7.5, 150 mM NaCl, and 0.1 % Tween 20) overnight at 4°C. All reagents used in these experiments were obtained from BioShop Canada unless otherwise indicated.

Blocked blots were washed three times in 1 × TBST washes before incubation with pre-chilled mouse anti-DnaK antibody (1:10,000 Ab in 3 % BSA and 0.01 % sodium azide in TBST; Enzo Life Sciences, Inc.) for one hour. This was followed by a second round of TBST washes before incubation with HRP conjugated goat anti-mouse Ab (1:5000 Ab in 3 % BSA in TBST; Promega Corp.) for an hour. After a third round of TBST washes, enhanced chemiluminescence (ECL) substrates A (1.5 mM Tris-HCl pH 8.8, 90 mM coumaric acid, and 250 mM luminol) and B (30 % H_2_O_2_ in dH_2_O) were mixed in a 1000:3 ratio and added to the blots prior to visualization via Chemidoc Touch Imaging system (BioRad). For OmpT visualization, blots were washed in TBST, and exposed to rabbit anti-OmpT antibody (1:10,000 Ab in 3% BSA and 0.01% sodium azide in TBST; Flare Biotech LLC) following by HRP conjugated goat anti-rabbit antibody (1:10,000 Ab in 3 % BSA in TBST; Cell Signaling Technology Inc). Images obtained were analyzed via ImageJ software.

### Construction of *ompT* promoter-omptin fusions

We used splicing by overlap extension polymerase chain reaction (SOE-PCR) to create fusions between the promoter of chromosomal ompT with the coding regions of ompT, ompP or arlC. These PCR products were then cloned into pCR2.1-TOPO according the manufacturer’s instructions. These constructs were then transformed to BL21 cells which lack a chromosomal *ompT* gene. Activity of the constructs were assessed by the FRET assay as described above.

## Results

### Omptin activity OmpT expression is regulated by limiting Mg^2+^

In order to determine how omptin protease activity is regulated we developed a FRET assay based on an internal fragment of the human host-defense peptide, LL-37. As seen in figure 1A, growth of BW25113 in N-minimal medium containing 10 μM Mg^2+^ results in ~9-fold induction of protease activity relative to growth in N-minimal medium with 2 mM Mg^2+^. In contrast, growth in the presence of 100 μM Fe^3+^ and 2 mM Mg^2+^ does not induce this activity. Finally, we examined the production of OmpT protein using western blotting. As seen in Figure 1B, low Mg^2+^, but not high Fe^3+^ resulted in ~6-fold upregulation of the OmpT signal relative to the control DnaK control.

**Figure 1:**
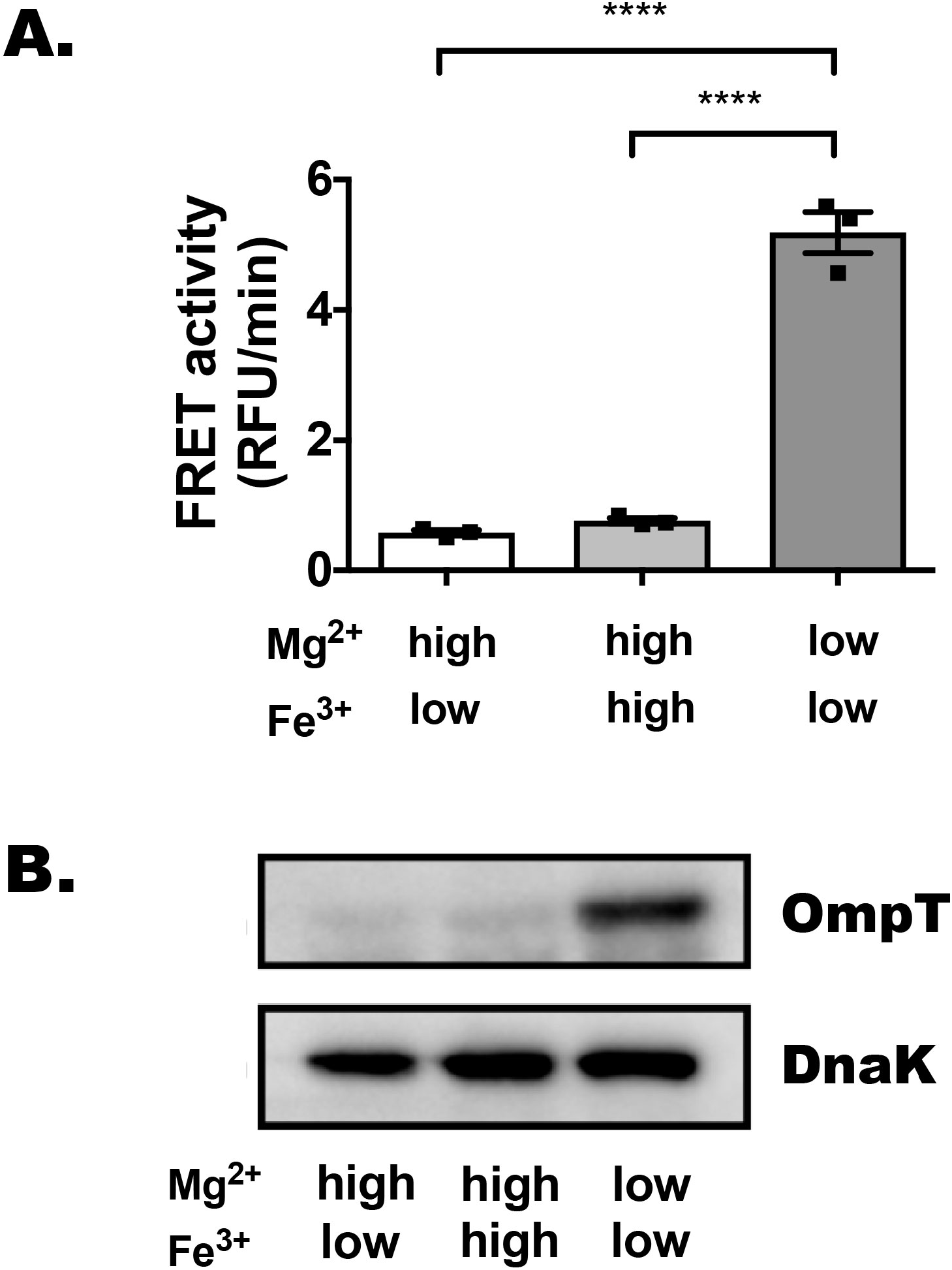
OmpT activity and expression are upregulated during growth in low Mg^2+^ conditions. A. Forster-resonance energy transfer (FRET) based activity of whole cell, mid-logarithmic phase culture of BW25113 grown in either 10 mM Mg^2+^, 1 µM Fe^3+^ N-minimal medium (high Mg^2+^, low Fe^3+^), 10 mM Mg^2+^ and 100 μM Fe^3+^ (high Mg^2+^, high Fe^3+^) or 10 μM Mg^2+^ and 1 μM Fe^3+^ (low Mg^2+^, low Fe^3+^) using an LL-37 based probe as described in the materials and methods. Results shown are the mean and standard error of the values represented by the filled squares indicated Statistics indicate p-values from one-way ANOVA NS – no statistical difference, * p<0.05, ** p<0.01, *** p<0.001, **** p<0.0001. B. Western blotting for OmpT proteins prepared from mid-logarithmic grown cultures under the conditions indicated.

### Expression and activity of *ompT* is regulated in a limiting Mg^2+^ and PhoP-dependent manner

The results above are consistent with PhoP-dependent and PmrA-independent regulation of OmpT expression and omptin protease activity. We examined FRET activity in BW25113 grown to mid-log growth (OD600 ~0.6) in BW25113 or isogenic ΔpmrA or ΔphoP mutants. As seen in Figure 2A, we observed a very significant reduction in FRET activity in the ΔphoP mutant. We wanted to examine whether the PhoP-dependent activity observed above was also present at the transcriptional level. We created plasmid-based *ompT*-GFP fusions and transformed these to BW25113, BW25113ΔphoP and BW25113ΔpmrA. We assessed transcriptional activity in N-minimal medium containing either 2 mM or 10 µM Mg^2+^. As observed in Figure 2B, low Mg^2+^ resulted in significant increases in GFP signal in both BW25113 and the ΔpmrA mutant. In contrast, deletion of *phoP* resulted in complete loss of low Mg^2+^ induced *ompT*-GFP expression. Consistent with this, loss of functional PhoP also resulted in almost complete loss of OmpT protein as assessed by Western blot (Figure 3C). Deletion of PmrA had no effect on OmpT expression.

**Figure 2.**
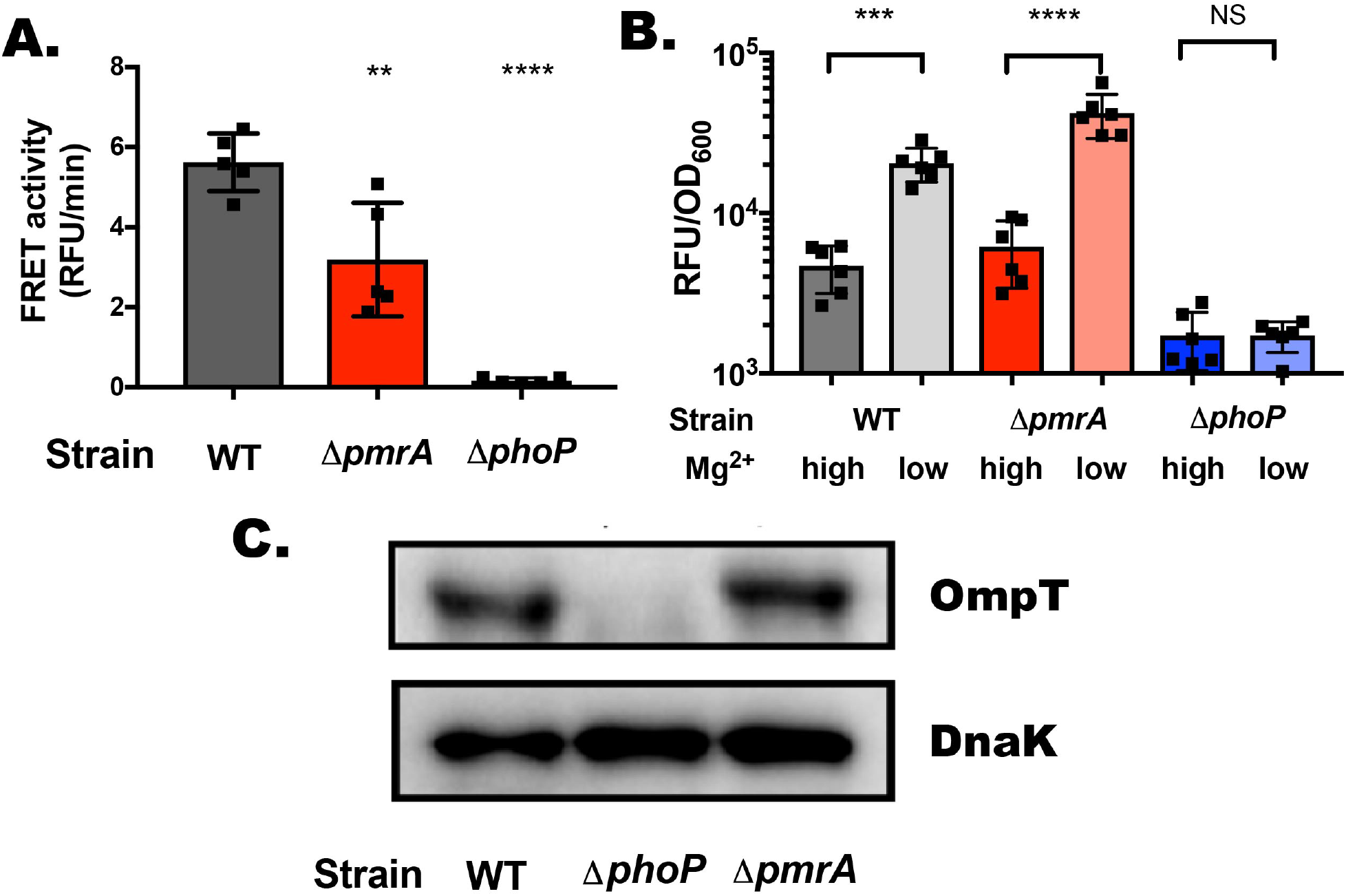
OmpT expression in BW25113 is regulated by low Mg2+ conditions in a PhoP-dependent manner. A. A. Forster-resonance energy transfer (FRET) based activity of whole cell, mid-logarithmic phase culture of BW25113, BW25113ΔpmrA and BW25113ΔphoP grown in LB. Results shown are the mean and standard error of the values represented by the filled squares indicated. Statistics indicate p-values from one-way ANOVA; NS – no statistical difference, * p<0.05, ** p<0.01, *** p<0.001, **** p<0.0001 B) Transcriptional activity of a plasmid-based ompT-GFP fusion protein in BW25113, BW25113ΔpmrA and BW25113ΔphoP grown in N-minimal medium containing 10 mM (high) Mg2+ or 10 µM (low) Mg2+. Results shown are the mean and standard error of the values represented by the filled squares indicated C) Western blotting for OmpT proteins prepared from mid-logarithmic grown cultures of BW25113, BW25113ΔpmrA and BW25113ΔphoP in N-minimal medium containing 10 μM Mg2+.

### Omptin activity in diverse enterobacteriaceal strains is also PhoP-dependent

In *E. coli*, three omptin proteins, OmpT, OmpP and ArlC have been described to date and these have been linked to degradation of cationic molecules in the bacterial environment. In addition, promoter diversity in the *ompT* gene has also been associated with elevated omptin production in enterohemorrhagic *E. coli* O157:H7 (Thomassin *et al.*, 2012). *Salmonella enterica* and *Citrobacter rodentium* are other Enterobacteriaceae that encode an OmpT orthologue (PgtE and CroP respectively). In order to test the PhoP-dependence of omptin activity in these strains, we obtained isogenic PhoP or PhoPQ mutants of each strain and subjected the strains to FRET assay. As seen in Figure 3, deletion of the *phoP* gene from three additional *E. coli* strains: AIEC strain LF82 (Boudeau *et al.*, 1999; McPhee *et al.*, 2014), AIEC strain NRG857c (Eaves-Pyles *et al.*, 2008; McPhee *et al.*, 2014), or EHEC strain 86-24 (Donnenberg *et al.*, 1993; Kunwar and Foster, unpublished) also abrogated omptin activity in these strains. We also obtained a PhoPQ mutant in *Citrobacter rodentium* (Reid-Yu *et al.*, 2015). Consistent with what we observed in all of the *E. coli* strains, we also observed complete abrogation of omptin protease activity in these strains when PhoP is absent.

**Figure 3.**
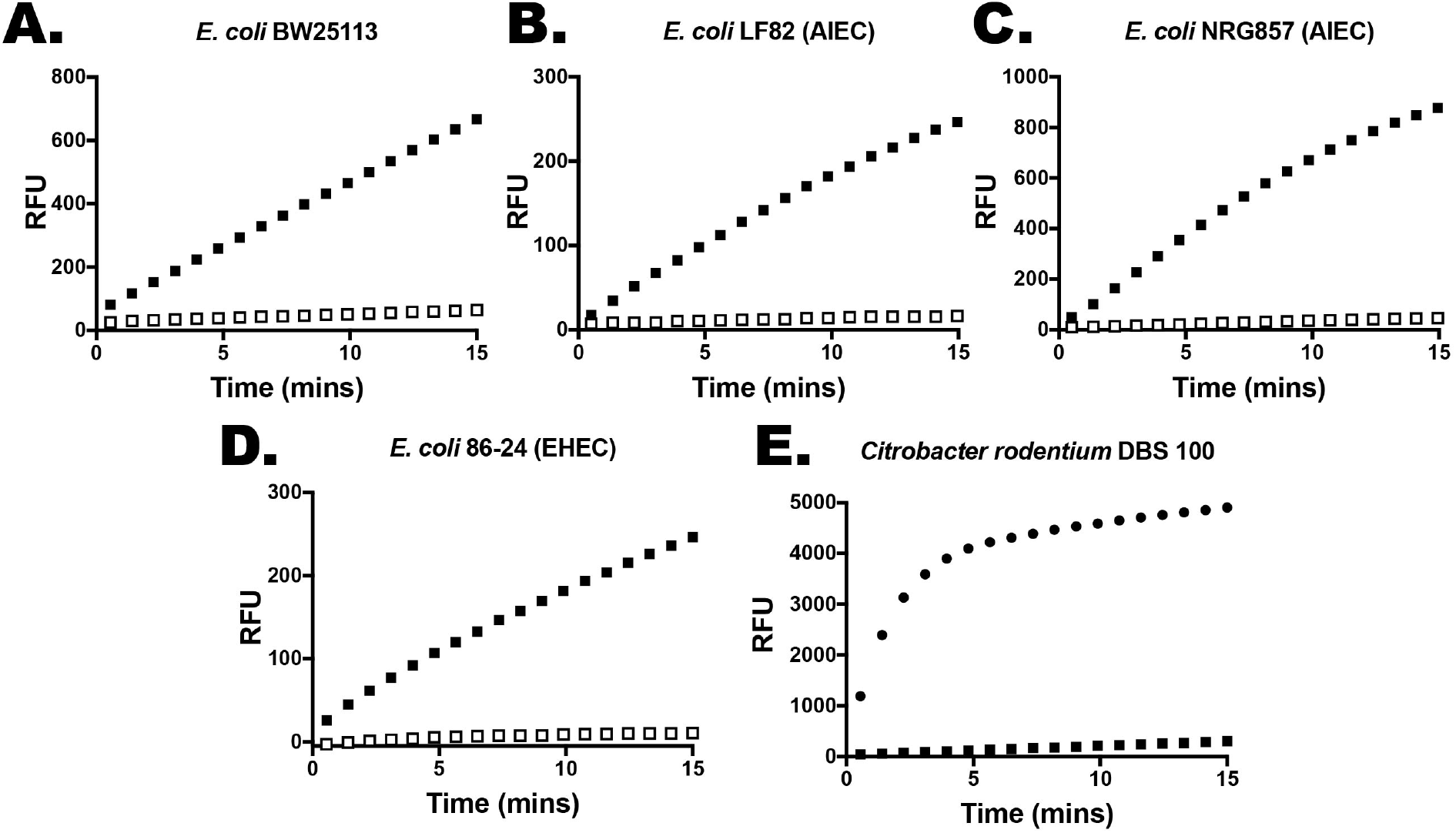
Divergent strains of Enterobacteriaceae exhibit PhoP-dependent omptin activity. Forster-resonance energy transfer (FRET) based activity of whole cell, mid-logarithmic phase cultures of either WT (black boxes) or ΔphoP (white boxes) strains of each of the bacterial strains indicated. Results shown are the mean value of three independent biological replicates of the samples as indicated.

### PhoP-dependent regulation of OmpT orthologues, OmpP, ArlC, and CroP

Our observation that omptin activity in strains/species containing diverse omptin orthologues in multiple species of Enterobacteriaceae led us to test whether the transcriptional regulation observed for ompT-GFP was also conserved in these other strains/proteins. We constructed a series of GFP reporter fusions, including *ompP*-GFP, *arlC*-GFP, and *croP*-GFP. These plasmid-encoded fusions were then transformed to appropriate host strains, including *E. coli* BW25113 or BW25113ΔphoP, and *C. rodentium* DBS100 or DBS100ΔphoPQ. As seen in figure 4, fluorescence from these constructs was controlled in a PhoP-dependent manner.

**Figure 4.**
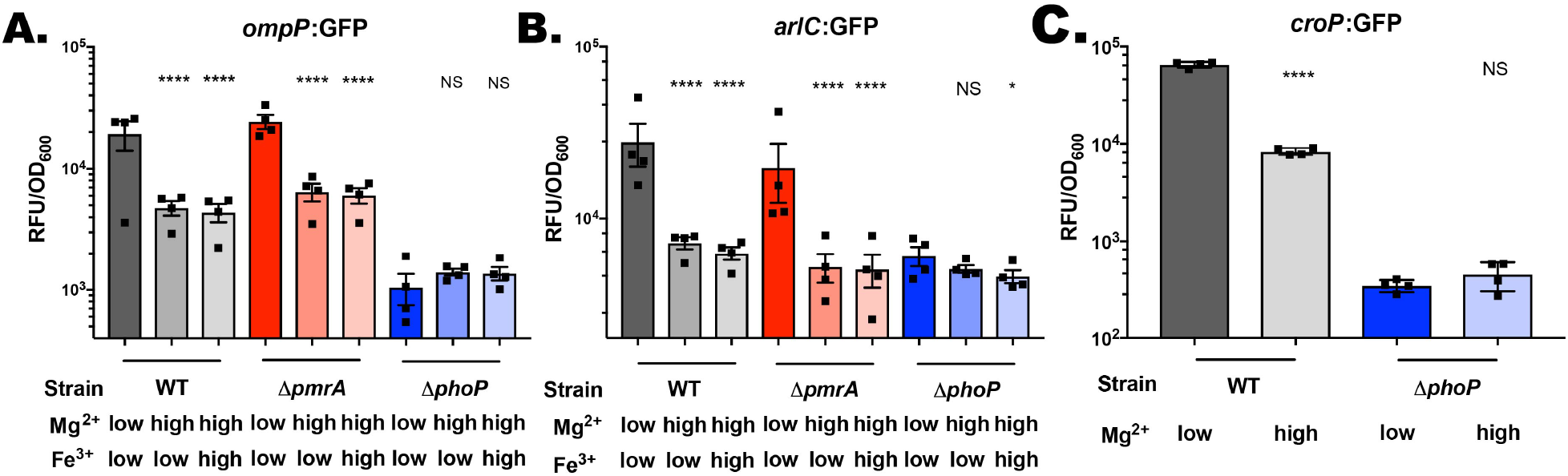
Diverse omptin orthologues are transcriptionally regulated by limiting Mg2+ in a PhoP-dependent manner. A plasmid-based omptin:GFP transcriptional reporter fusion was transformed into BW25113, BW25113ΔpmrA and BW25113ΔphoP and the fluorescence was examined during growth in N-minimal medium containing 10 mM Mg2+ and 1 μM Fe3+ (high Mg2+, low Fe3+), 10 mM Mg2+ and 100 μM Fe3+ (high Mg2+, high Fe3+) or 10 μM Mg2+ and 1 μM Fe3+ (low Mg2+, low Fe3+) as indicated in the figure. A) *ompP*:GFP fusion. B) *arlC*:GFP fusion. C) *croP*:GFP fusion. Results shown are the mean and standard error of the values represented by the filled squares indicated. Statistics indicate p-values from one-way ANOVA NS – no statistical difference, * p<0.05, ** p<0.01, *** p<0.001, **** p<0.0001

### PhoP dependent regulation of OmpT requires a conserved PhoP-DNA binding site

Genes that are directly PhoP-regulated usually contain a specific PhoP-binding motif, termed the PhoP-box. We looked for a PhoP-box in the promoters of OmpT, OmpP and ArlC of *E. coli* as well as CroP of *C. rodentium* and identified a strongly conserved site on the reverse strand that closely matched the published consensus sequence identified in Zwir et al, 2010 (Figure 5A). We made an AAA → GGG mutation in the most gene-adjacent half of the DNA binding site in our *ompT*-GFP reporter fusion (Figure 5B). As seen in figure 5C, this mutation resulted in a complete loss of low Mg^2+^ and PhoP dependent induction of *ompT*-GFP activity, indicating that this PhoP-binding site was critical for low-Mg2+ induced *ompT* transcriptional activity.

**Figure 5.**
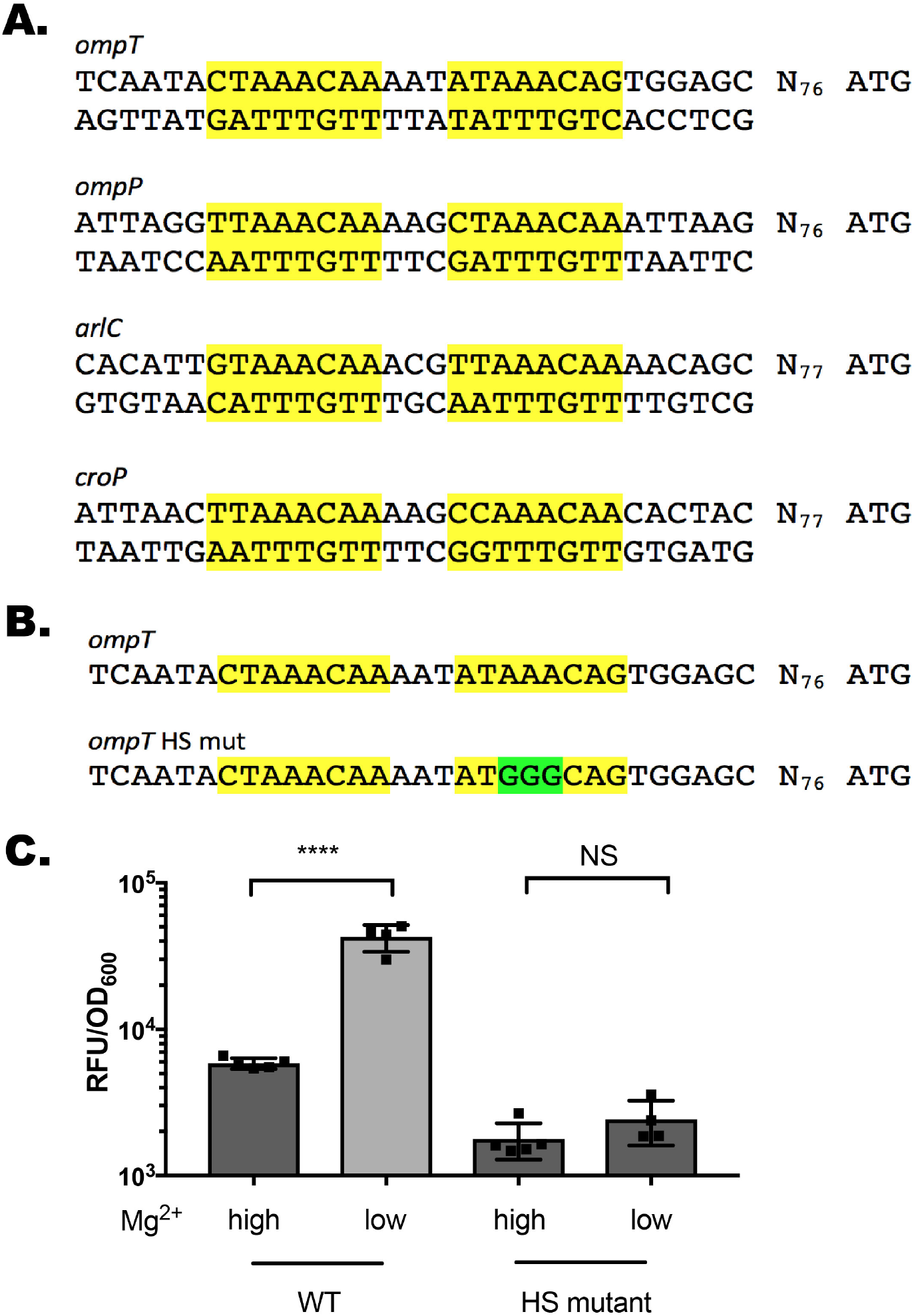
Identification of a PhoP-binding site required for low Mg^2+^ regulated *ompT*:GFP fusion activity. A) Identification of conserved putative PhoP-binding sites in omptin family proteins of *Escherichia coli* (*ompT*, *ompP*, and *arlC*) and *Citrobacter rodentium* (*croP*). B) Sequence of the engineered AAA → GGG half-site mutation in the *ompT*:GFP promoter. C) Fluorescence of *ompT*:GFP or *ompT*HSmut:GFP fusions in N-minimal medium containing 10 mM Mg^2+^ (high) or 10 μM Mg2+ (low Mg^2+^) as indicated in the figure. Results shown are the mean and standard error of the values represented by the filled squares indicated. Statistics indicate p-values from one-way ANOVA NS – no statistical difference, * p<0.05, ** p<0.01, *** p<0.001, **** p<0.0001

### Omptin orthologues of *E. coli* exhibit differential activity toward specific substrates

Although all regulated by the same transcriptional regulator, it is possible that observed differences in activity could result from different degrees of induction due to subtle promoter differences between omptin orthologues. We created plasmid based fusions in which we put the genes for each *E. coli* omptin orthologue (*ompT, ompP, arlC*) under the control of the *ompT* promoter in order to study the activity of the individual proteins alone independent of cognate promoter strength. These fusions were then transformed to the omptin-deficient strain BL21. We then carried out a FRET reporter assay on these transformed strains. As shown in figure 6, we observe that despite being controlled by the same promoter, there are significant differences in the activity of each fusion toward this substrate, suggesting that each protease has differential activity/selectivity toward different protease substrates.

**Figure 6.**
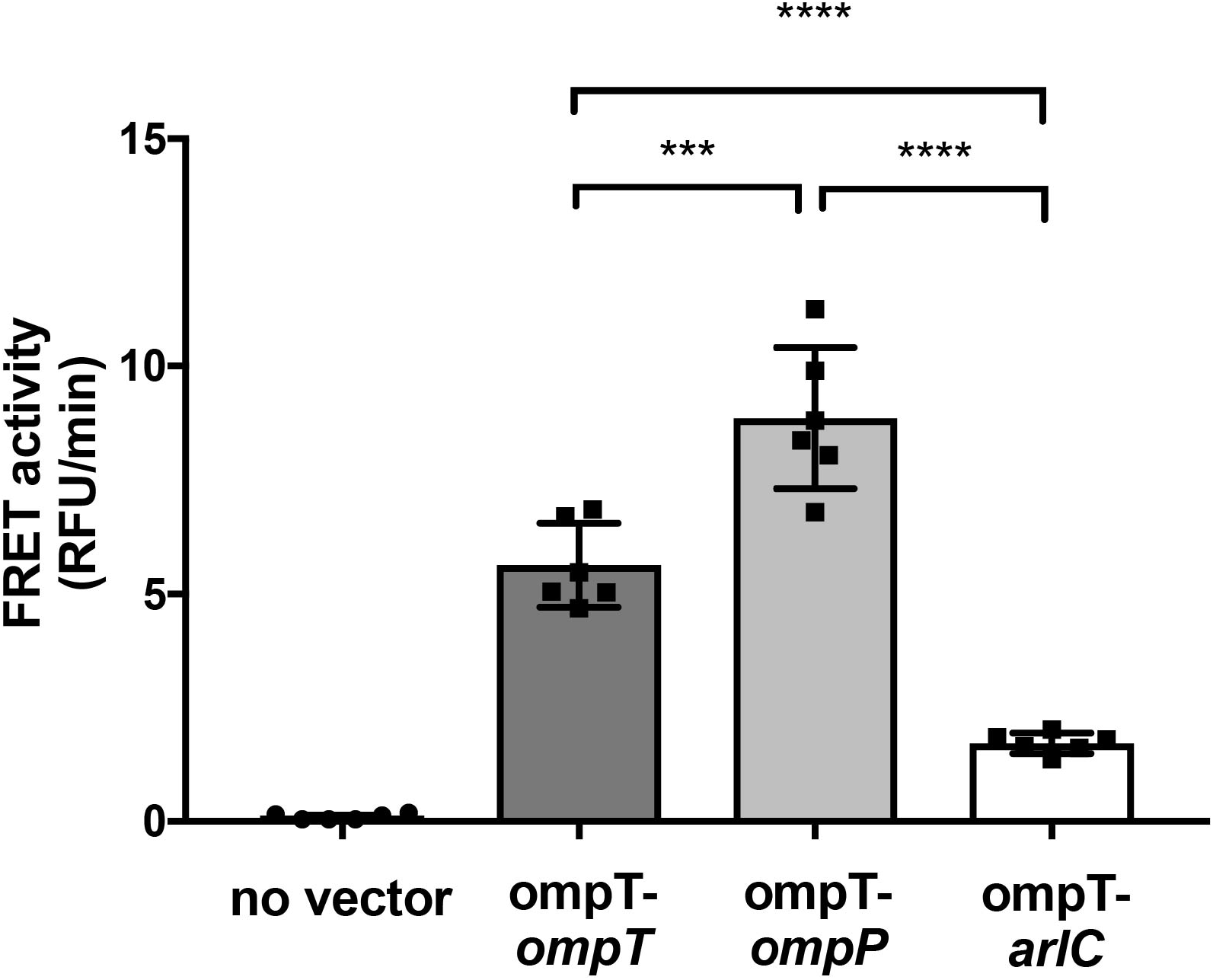
Differential activity of *ompT* promoter-omptin gene fusions to an LL-37 based FRET activity probe. The *ompT* promoter-omptin gene fusion constructs were transformed into the omptin deficient strain BL21 and assessed for protease activity via FRET assay at mid-logarithmic phase. Results shown are the mean and standard error of the values represented by the filled squares indicated. Statistics indicate p-values from one-way ANOVA NS – no statistical difference, * p<0.05, ** p<0.01, *** p<0.001, **** p<0.0001

## Discussion

Bacteria have evolved numerous strategies to overcome antimicrobial pressure experienced during host infection. Among the most important regulators of virulence and host-mediated antimicrobial defense is the PhoPQ two-component regulatory system. The PhoPQ system responds to elevated concentrations of cationic host-defense peptides (Bader *et al.*, 2005), to reduced pH (Prost and Miller, 2008), to changes in osmolarity (Yuan *et al.*, 2017) and to limiting Mg^2+^ conditions (García Véscovi *et al.*, 1996). Similarly, the PmrAB system responds to high concentrations of ferric iron and low pH (Wösten *et al.*, 2000). Following activation of the membrane-bound sensor kinase by any of the previously mentioned signals, each response regulator directly regulates a subset of genes. Upon phosphorylation, PhoP binds to DNA motifs in the promoters of PhoP-regulated genes and activates their transcription (Zwir et al, PNAS 2005). One of the PhoP-regulated genes, PmrD, binds to phospho-PmrA and prevents its dephosphorylation (Kox *et al.*, 2000; Kato and Groisman, 2004). This results in PhoP-mediated control of PmrA-regulated genes (Rubin *et al.*, 2015). This is important, because PmrA regulates a number of genes that change the structure and charge of LPS. The PhoPQ system is crucial for virulence in many enteric pathogens, including *Salmonella enterica*, *E. coli* and *Yersinia* sp. (Miller *et al.*, 1989; Grabenstein *et al.*, 2004; Alteri *et al.*, 2011).

Activation of the PhoPQ system is well-established as a mechanism for the regulation of outer membrane susceptibility to cationic antimicrobial peptides (Dalebroux and Miller, 2014). Much of this resistance occurs via modifications to the structure of lipopolysaccharide that alter the surface charge and amphipathicity of the molecule, leading to reduced interaction with cationic antimicrobial peptides. These modifications include the addition of a palmitoyl residue to an oxyacyl site in lipid A via the PagP protein, and the masking of surface anionic charge via the addition of 4-aminoarabinose and/or ethanolamine to the 1 and/or 4’ phosphates of lipid A (Bishop *et al.*, 2000; Trent *et al.*, 2001; Lee *et al.*, 2004). Other modifications are possible including the masking of other phosphates in the LPS core or dephosphorylation of lipid A or core (Chen and Groisman, 2013). The net effect of these modifications, whatever the specific changes made, are resistance to cationic host defense peptides.

In addition to this LPS-modification mediated antimicrobial peptide resistance, enterobacteriaceae also produce omptin proteases that can target components of the host innate immune system. Omptins are 10-stranded β-barrel proteins that traverse the bacterial outer membrane. Crystal structures of the prototypical omptin, OmpT, show that the surface loops of the barrel protrudes ~40 Å above the lipid bilayer into the extracellular space and the catalytic site of the protease is contained within the extracellular loops (Vandeputte-Rutten *et al.*, 2001). The catalytic site for omptins includes a strictly conserved His-Asp dyad that activate a nucleophilic water molecule and an Asp-Asp dyad that is believed to stabilize the intermediate state (Eren *et al.*, 2010). Omptin proteases exhibit a strong preference for cleavage at dibasic sites, although some exceptions to this exist (Brannon *et al.*, 2015). This preference appears to be due to a fairly anionic pocket at the base of the loop region and adjacent to the catalytic site (Vandeputte-Rutten *et al.*, 2001). As with many other β-barrel/porin family proteins, omptins are tightly associated with bacterial LPS, can form co-crystals with LPS, and require LPS for functional activity (Kramer *et al.*, 2002; Eren and van den Berg, 2012).

Omptin proteases have a wide variety of substrates that may contribute to in vivo fitness of the bacterium containing them, including HDPs like LL-37 or CRAMP (McPhee *et al.*, 2014; Brannon *et al.*, 2015), components of the complement cascade or the clotting cascade (Lathem *et al.*, 2007; Riva *et al.*, 2015), and non-physiological but cationic substrates like protamine (Johanna Haiko *et al.*, 2009). Alterations in either the activity of omptins or in their transcriptional activity have been associated with changes in virulence, including in pandemic *Y. pestis* (Haiko *et al.*, 2009) or in invasive *Salmonella enterica* (Hammarlöf *et al.*, 2018) suggesting that omptin proteases are critical factors for interaction with and resistance to components of the host innate immune system and that understanding both the mechanism(s) by which they are regulated and the substrates with which each omptin interacts may shed light on their role in pathogenesis.

In *E. coli*, at least three omptin proteases have been identified to date: the housekeeping chromosomally-encoded OmpT (Grodberg and Dunn, 1988), the F’ episome encoded OmpP (Kaufmann *et al.*, 1994) and a plasmid encoded protein, ArlC (McPhee *et al.*, 2014). *Citrobacter rodentium* encodes the OmpT orthologue CroP (Le Sage *et al.*, 2009). Although these proteins have been associated with in vivo fitness and virulence, a detailed examination of their mechanism of regulation has not been undertaken. Here, we show that the housekeeping omptin OmpT is regulated by the PhoP transcriptional regulator, consistent with previous data (Eguchi *et al.*, 2004). We identified a putative PhoP-binding site in the promoter of OmpT and we show that mutation of this site abrogates PhoP-dependent regulation. We also identified similar putative PhoP-binding sites in the promoters of *ompP*, *arlC* and *croP* and clearly demonstrate that these omptins are also PhoP-regulated.

Although we show here that PhoP regulation of the *E. coli* omptins is conserved, there may be alterations in the promoter sequence that could affect the expression levels of the protein, with significant impacts on virulence. This has been demonstrated previously in enterohemorrhagic *E. coli*, in which the *ompT* promoter in EHEC supports significantly increased expression of OmpT protein, with increased resistance to LL-37 (Thomassin *et al.*, 2012). In addition, other regulatory inputs could contribute to altered expression levels among the omptin orthologues could alter activity of these proteins. This has been demonstrated previously for the PgtE protease of *Salmonella*, which is subject to significant silencing by Hns (Will *et al.*, 2014) and positive regulation by the SlyA and PhoP protein and (Navarre *et al.*, 2005). Even the prototypical OmpT protein shows evidence of regulation by small RNAs (Guillier and Gottesman, 2006). Our data here suggests that regardless of the increased activity observed in EHEC OmpT, the activity the *E. coli* and *C. rodentium* omptins is wholly dependent on PhoP. Whether other environmental conditions or regulatory proteins might further contribute to differential omptin regulation remains an unanswered, but important, question.

The observation that omptin regulation by PhoP is conserved in OmpT, OmpP, ArlC and CroP raises an interesting question, “why would closely related bacteria maintain equivalent regulatory control over multiple different omptin alleles”? We hypothesized that each allele might provide substrate selectivity that might favour the maintenance of one allele or another, depending upon the strain and the environments in which that strain is found. Divergent omptin substrate selectivity has been observed previously between OmpT and CroP, in which OmpT preferentially cleaves antimicrobial peptides with α-helical structure while CroP exhibits a preference for unfolded/unstructured peptides (Brannon *et al.*, 2015). Similarly, the *Shigella* virulence-related omptin protein, SopA has lost generic broad-substrate activity and appears to have become tightly tuned to a single substrate (IcsA) and maintains this substrate with a strict polar localization (Egile *et al.*, 1997; Shere *et al.*, 1997). This suggests that the multiple *E. coli* omptin orthologues described to date may also exhibit altered substrate specificity.

Comparison of OmpP and OmpT have identified some differences in the substrate specificity of the enzymes, although both were able to protect the expressing strains from protamine exposure (Hwang *et al.*, 2007). Similarly, previous work has shown that both OmpT and ArlC contribute to host-defense peptide resistance, with significant differences in the resistance profile of isogenic strains lacking either ArlC or OmpT, suggesting that these proteins have different substrate specificities (McPhee *et al.*, 2014). These observations bear that caveat that the proteins studied were under the control of the native promoter and it is therefore difficult to distinguish between altered expression levels and altered substrate specificity. We attempted to remedy this caveat by expressing each of the *ompT*, *ompP* and *arlC* genes from the *ompT* promoter. As expected, this demonstrated that each omptin had significantly different activity from one another, supporting the hypothesis that each omptin may have preferred substrates in vivo.

Understanding the mechanisms for by which omptin proteins are regulated as well as how they differ in their preferred substrate may shed light on how virulence emerges in these microbes. Although OmpT appears to be conserved in the majority of *E. coli* strains, both OmpP and ArlC are usually associated with mobile genetic elements, suggesting that the acquisition of one of these proteins could open up new niches for colonization or invasion by these microbes. Here, we provide evidence that although the omptins of *E. coli* are regulated in a conserved manner, they still exhibit significant differences in behaviour – supporting the hypothesis that they might be associated with altered virulence or host adaptation.

## Acknowledgements

This work was supported by a National Sciences and Engineering Research Council Discovery Grant (RGPIN-04679-2015) and by startup funding from the Ryerson University Faculty of Science to JBM.

